# Inhibitory cell type heterogeneity in a spatially structured mean-field model of V1

**DOI:** 10.1101/2025.03.13.643046

**Authors:** Soon Ho Kim, Hannah Choi

## Abstract

Inhibitory interneurons in the cortex are classified into cell types differing in their morphology, electrophysiology, and connectivity. Although it is known that parvalbumin (PV), somatostatin (SST), and vasoactive intestinal polypeptide-expressing neurons (VIP), the major inhibitory neuron subtypes in the cortex, have distinct modulatory effects on excitatory neurons, how heterogeneous spatial connectivity properties relate to network computations is not well understood. Here, we study the implications of heterogeneous inhibitory neurons on the dynamics and computations of spatially-structured neural networks. We develop a mean-field model of the system in order to systematically examine excitation-inhibition balance, dynamical stability, and cell-type specific gain modulations. The model incorporates three inhibitory cell types and excitatory neurons with distinct connectivity probabilities and recent evidence of long-range spatial projections of SST neurons. Position-dependent firing rate predictions are validated against simulations, and balanced solutions under Gaussian assumptions are derived from scaling arguments. Stability analysis shows that while long-range inhibitory projections in E-I circuits with a homogeneous inhibitory population result in instability, the heterogeneous network maintains stability with long-range SST projections. This suggests that a mixture of short and long-range inhibitions may be key to providing diverse computations while maintaining stability. We further find that conductance-based synaptic transmissions are necessary to reproduce experimentally observed cell-type-specific gain modulations of inhibition by PV and SST neurons. These gain modulations are distance-dependent, and a linear response analysis suggested that shifts in the excitation-inhibition balance of the network underlie these effects. Our theoretical approach offers insight into the computational function of cell-type-specific and distance-dependent network structure.

## I. INTRODUCTION

Cortical processing involves a precise interplay between synaptic excitation and inhibition [1–3]. Inhibitory neurons tune cortical oscillations, modulate gain, and gate excitatory neuronal responses [4–7]. Although cortical circuits are commonly modeled as recurrent neural networks [8], biological circuits have two features often overlooked in model networks: spatially-dependent connectivity [9] and the existence of cell types [10, 11].

Biological neural networks are spatially embedded, and distance plays an important role in governing connection probabilities [12]. To describe their effects on network dynamics, mean-field models of spatially distributed spiking neural networks have been developed in previous studies [13–15]. These studies have found that broader lateral projections can result in distinct pairwise correlation structures [15]. Furthermore, stability analysis of these models shows that the distance of projections has a critical effect on network stability: if inhibitory projections are broader than the excitatory projections, the fixed points of the network can become unstable [13], which can result in unreliable tracking of inputs [14]. These studies, however, have not examined the consequences of having cell types and assumed homogeneous inhibitory populations.

In cortical networks, diverse inhibitory subtypes including parvalbumin (PV), somatostatin (SST), and vasoactive intestinal polypeptide-expressing neurons (VIP) form circuits with excitatory (E) neurons with distinct connectivity statistics [10, 16, 17]. Several theoretical works have examined networks with heterogeneous inhibitory cell types. Wilson-Cowan-type models incorporating cell types have been used to study inhibitory stabilization and subtractive/divisive modulations [18] and to describe visual processing of objects superimposed on a background [19]. A rate-based circuit model has shown that PV and SST neurons may each be involved in encoding the mean and variance of predictions in prediction error neurons [20]. Recent work has studied the interplay between the inhibitory pathway from SST to E and the disinhibitory pathway involving PV, showing that SST can exhibit multiple gain modulations depending on the synaptic weights. [21] These works have suggested the role of cell types in network computations, but did not address realistic, spatially-structured connectivity among different cell types and in particular did not address distance-dependent modulations.

Understanding how cell types and their specific distance-dependent connectivity influences neural computations is an important aspect of cortical processing that is not well understood. The computational functions of inhibitory neurons can be understood through their arithmetic operations on neuronal input-output functions [22]. In the mouse primary visual cortex (V1), local and moderate activation of PV neurons results in divisive modulations on excitatory response curves to stimuli while the effect of SST neurons is subtractive [23– 27]. However, recent evidence shows that when activated distally, SST neurons inhibit excitatory neurons receiving visual stimuli in a divisive manner while PV neurons exert subtractive modulations [28]. Anatomical data showed that SST neurons form broader axonal projections than PV neurons do [28–30]. Furthermore, a computational model of spiking neurons suggested that this long-range inhibition is necessary to reproduce the effects [28]. On the other hand, VIP neurons may increase the response gain, possibly through disinhibition [7, 31].

While spatially-embedded spiking neural networks of cortical populations incorporating cell types have recently been developed [28, 32], the nature of large spiking network models has limited these previous studies to simulation-based analyses, lacking theoretical insights. A systematic analysis of how position-dependent activation of PV, SST and VIP neurons modulates neural activity, therefore, is missing. A biologically-informed computational model of V1 enabling such analysis *in silico* would provide mechanistic insight into these experimental results. A mean-field model in particular can be tractable under some assumptions, allowing direct computation of important network features such as the eigenspectrum of the linearized dynamics. Furthermore, such models would provide a platform to understand the role of celltype-specific structure in information processing across cortical networks.

Here we develop a mean-field model of spatially-extended spiking neural networks in V1. We first construct the model, incorporating distinct, cell-type-specific connectivity and derive conditions that achieve excitatory-inhibitory balance in the thermodynamic limit. We use an approach based on the Fokker-Planck equation to compute the position-dependent stationary firing rates of the model and validate them against spiking neural network simulations. We examine stability features of the network, specifically how long-range SST projections affect network stability. We further characterize the spatiotemporal features of the network dynamics across the stability boundary, finding significant differences in the heterogeneous network. Finally, we examine gain modulations during activation of specific inhibitory cell types by examining shifts in the stationary firing rates with input strength and find cell-type-specific and spatially-dependent effects. A linear response analysis suggests that shifts in the balance of excitation and inhibition through disexcitation or disinhibition may be a mechanism for the differential modulations.

## II. RESULTS

### A. Network Model with Inhibitory Cell Types

We construct a spatially-organized network of spiking neurons and incorporate cell-type-specific connectivity. The model consists of *N* neurons that are uniformly distributed on a 1-dimensional space with periodic boundary conditions, Γ = (0, 1] (i.e., a ring). The network consists of *N*_*e*_ = *q*_*e*_*N* excitatory and *N* − *N*_*e*_ inhibitory neurons which are subdivided into *N*_*p*_ = *q*_*p*_*N* PV, *N*_*s*_ = *q*_*s*_*N* SST, and *N*_*v*_ = *q*_*v*_*N* VIP neurons with *q*_*e*_ + *q*_*p*_ + *q*_*s*_ + *q*_*v*_ = 1. We set *q*_*e*_ = 0.8, *q*_*p*_ = 0.1, and *q*_*s*_ = *q*_*v*_ = 0.05, approximately following anatomical evidence [11] (see Figs. 1a, b for model schematics).

**FIG. 1.**
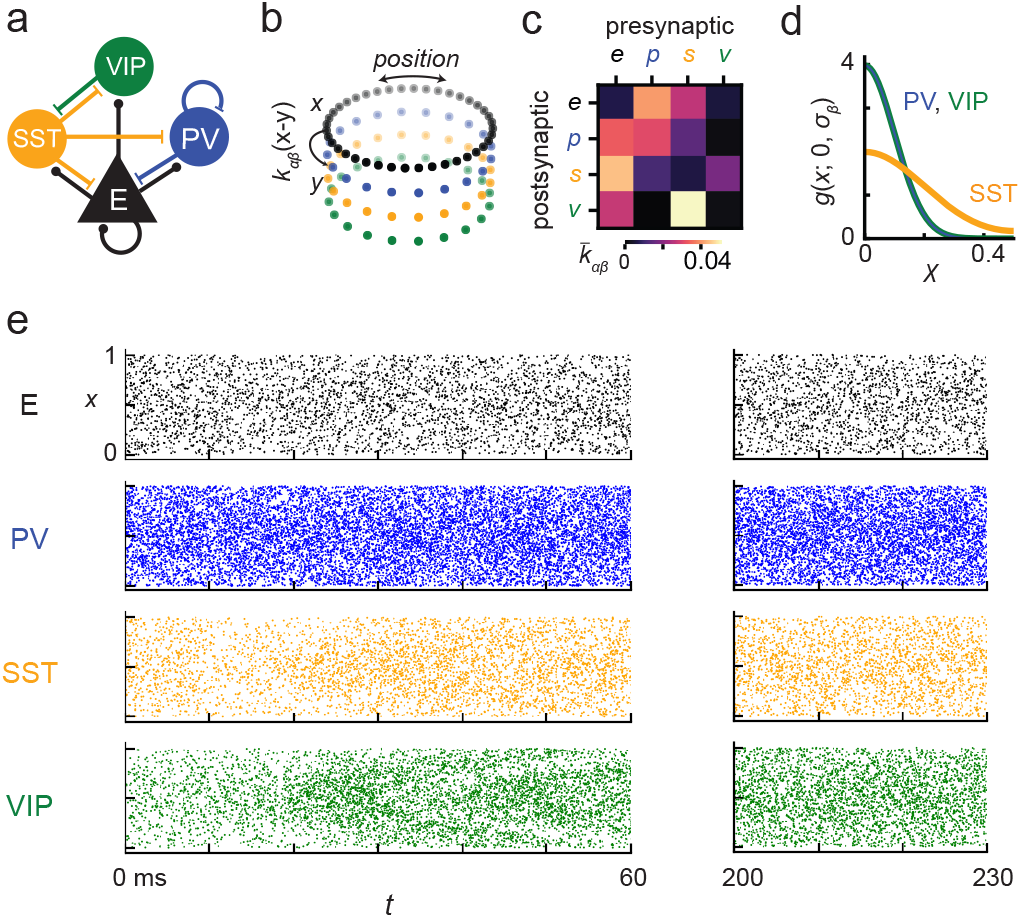
Model schematic and spiking dynamics. (a) The cortical network model consists of excitatory (E) neurons and PV, SST, and VIP inhibitory neurons. The schematic shows the major projections within and between cell types. (b) Cartoon of the spatial structure of the network. Neurons are placed with even spacing in a ring configuration. The probability of connection depends on the relative position *x − y* between the position of the presynaptic and postsynaptic neurons. (c) Cell-type-specific connectivity probabilities. Brighter color represents a greater probability that the corresponding presynaptic cell type will project to the postsynaptic cell type. Indices *e, p, s*, and *v* correspond to E, PV, SST, and VIP neurons, respectively. (d) Spatial projection kernel of each inhibitory cell type. (e) Raster plot of example spiking neural network run with a network size of *N* = 5 × 10^5^. The vertical axes represents the position and the horizontal axes the time of the spike.

We first assume that the neurons follow leaky integrate-and-fire (LIF) dynamics with current-based synapses (see Eqs. 9, 10 in Sec. IV A). The probability with which the *j*th neuron of cell type *β* at position *y* = *j/N*_*β*_ (where *β* takes on indices *e, p, s*, and *v* for E, PV, SST, and VIP neurons) connects to the *k*th neuron of type *α* at *x* = *k/N*_*α*_ is given by

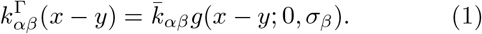

Here 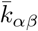 controls the overall connectivity probability from *β* to *α* (we use 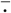 to denote the spatial average, i.e., 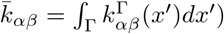 .We select relative values to match recent anatomical data from layer 2/3 of V1 in mice [17] (Fig. 1c; see also Table I in Methods). Additionally, we scale all probabilities so that the overall connectivity of the network is 1%, imposing sparse connectivity. The distance-dependence is given by the wrapped Gaussian function *g*(*x* − *y*; 0, *σ*_*β*_), where *σ*_*β*_ is the characteristic projection distance of *β*-type neurons (see Methods A). The values for *σ*_*β*_ are also chosen based on recent anatomical evidence; motivated by the experimental evidence of broad SST axonal projections and more local PV and VIP projections [28–30, 33], we set *σ*_*s*_ = 0.2 and *σ*_*p*_ = *σ*_*v*_ = 0.1 (Fig 1d). Excitatory projections are set to *σ*_*e*_ = 0.1 unless stated otherwise. If a synapse is formed from a presynaptic neuron of type *β* to a postsynaptic one of type *α*, the weight of the synapse is given by *J*_*αβ*_, where the synaptic weight matrix *J* is tuned to achieve a balanced state as outlined below.

**TABLE I.**
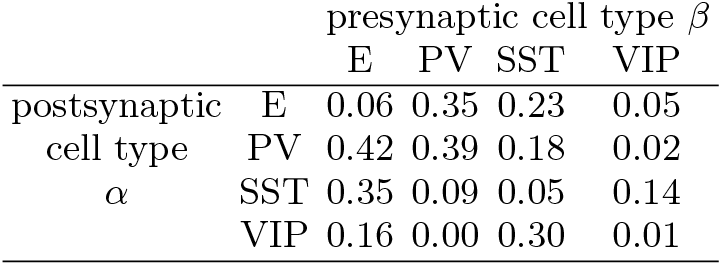
Cell-type-specific connectivity probabilities 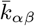 used in the model.

### B. Derivation of balanced solution in the thermodynamic limit

The *k*th neuron of cell type *α* receives inputs *I*_*α,k*_ given by a sum of recurrent synaptic inputs, described above, and a cell-type- and position-dependent external current *F*_*α*_(*x*). We consider a case where *F*_*α*_(*x*) have Gaussian spatial profiles. The external input to population *α* is given by a weighted sum of a constant term and spatially structured term, 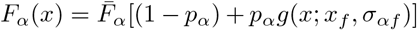, where the stimulus is centered at *x*_*f*_ = 0.5 and has a broadness given by *σ*_*αf*_ = 0.25. The relative weight of the spatially-structured input is given by *p*_*α*_, and 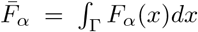 is the mean current input to population *α*. Following studies of networks with E-I balance [8, 13], we assume that the synaptic weights follow scaling 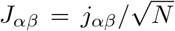 and inputs follow 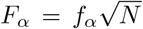 with network size *N* , where *j*_*αβ*_ and *f*_*α*_ do not depend on *N* . We assume that other parameters, such as the threshold and reset potentials, do not depend on *N* . Under these assumptions, a neuron receives *O*(*N*) excitatory inputs but requires 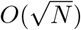 presynaptic neurons to activate during an integration window to produce a spike [34]. In order for a stable stationary firing rate solution to exist, a dynamic balance between excitation and inhibition must be maintained [8, 13, 35].

The *k*th neuron of type *α* (at position *x* = *k/N*_*α*_) has mean firing rate *ν*_*α*_(*x*) = ⟨*E*[*s*_*α,k*_]⟩, where *E*[·] denotes the average over network configurations, ⟨·⟩ the time average, and 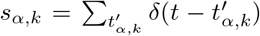 is the spike train of the neuron. The mean input current is given by

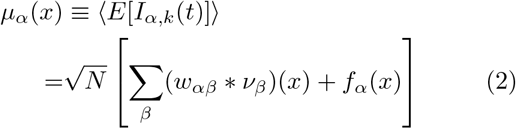

where 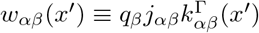 and “∗” indicates circular convolution, i.e., (*w*_*αβ*_ ∗*ν*_*β*_)(*x*) = ∫_Γ_ *w*_*αβ*_(*x*′)*ν*_*β*_(*x*−*x*′)*dx*′. The balanced state exists when the mean input currents obey 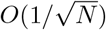 scaling,

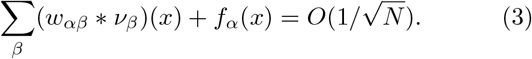

Taking *N* → ∞ (which we call the thermodynamic limit) gives a Fredholm equation of the first kind, which in the Fourier domain becomes

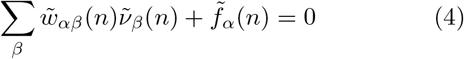

where the tilde indicates a Fourier transform, i.e. 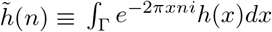, and *n* is the Fourier mode. The solution is given by

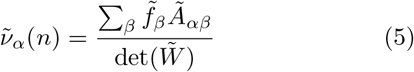

where 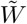 is a matrix whose elements are 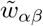 and *A*_*αβ*_ are cofactors of 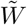. Eq. 5 must hold for every *n* for which det 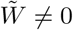. If the determinant equals 0 at some Fourier mode, then for a solution to exist, 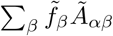 must also equal zero at that Fourier mode.

The solutions are viable only if 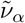 has a well-defined inverse Fourier transform, such that

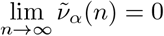

for each *α* = *e, p, s, v*. Each term (*w*_*αβ*_ ∗ *ν*_*β*_)(*x*) in Eq. 2 is a convolution of two Gaussian functions, which is itself a Gaussian with spatial variance equal to the sum of those of *w*_*αβ*_ and *ν*_*β*_. This implies that the external current inputs to each population *α* must be broader than the recurrent projections of every other population *β* (i.e., *σ*_*β*_ *< σ*_*αf*_ ∀*α, β*) to satisfy Eq. 4 [13]. Under these conditions, the stationary firing rates can be derived by taking the inverse Fourier transform of Eq. 5 to get (see Sec. IV C for details)

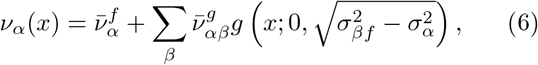

where 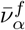 and 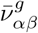 are defined in Eqs. 19-21.

If each input has the same profile, i.e. *σ*_*βf*_ = *σ*_*f*_ and *p*_*α*_ = *p*, Eq. 6 simplifies to

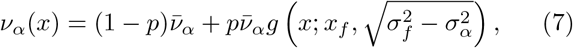

where 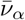 is the spatial mean of the firing rate. Synaptic weights *j*_*αβ*_ and input values 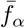 that achieve network balance can be generated through numerical means (see Sec. IV C and Table II). Spiking neural network simulations using balanced parameters are shown in Fig. 1e.

**TABLE II.**
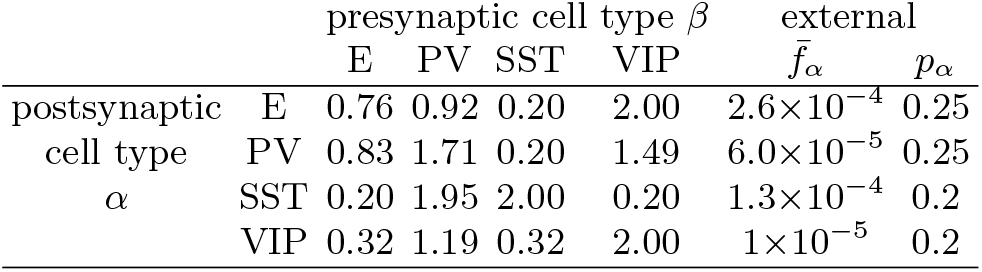
Synaptic weights *j*_*αβ*_ and external current input parameters 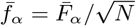 and *p*_*α*_ (see Eq. 13) used in Figs. 1-4. Values of *j*_*αβ*_ and 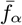 were chosen to achieve balance in the thermodynamic limit as described in the Methods.

### C. Finite-*N* solutions

Under mean-field assumptions, firing rates for networks of finite size *N* can be approximated with a semi-analytic approach. The current input *I*_*α,k*_ is replaced by a diffusion term 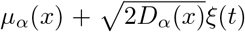 where *µ*_*α*_(*x*) is the mean and *D*_*α*_(*x*) the variance of the input to neurons of type *α*, and *ξ*(*t*) is a zero-mean white noise. Given *µ*_*α*_(*x*) and *D*_*α*_(*x*), the stationary firing rates *ν*_*α*_(*x*) can be computed by numerically integrating the Fokker-Planck equation as introduced in previous works [36, 37]. Since *µ*_*α*_(*x*) and *D*_*α*_(*x*) themselves depend on *ν*_*α*_(*x*), this forms self-consistent equations from which an iterative approach can be used to numerically compute the stationary firing rates (details in Sec. IV B).

Comparison of the stationary firing rates with LIF network simulations shows that the mean-field description accurately predicts the stationary firing rates (Fig. 2a). The normalized mean square error (NMSE) between the mean-field fixed points and rates computed from simulations decreases with network size (Fig. 2b). Furthermore, we see that as *N* increases the network converges to the analytical solutions satisfying the balanced state in the thermodynamic limit in Eq. 7 (Fig. 2c).

**FIG. 2.**
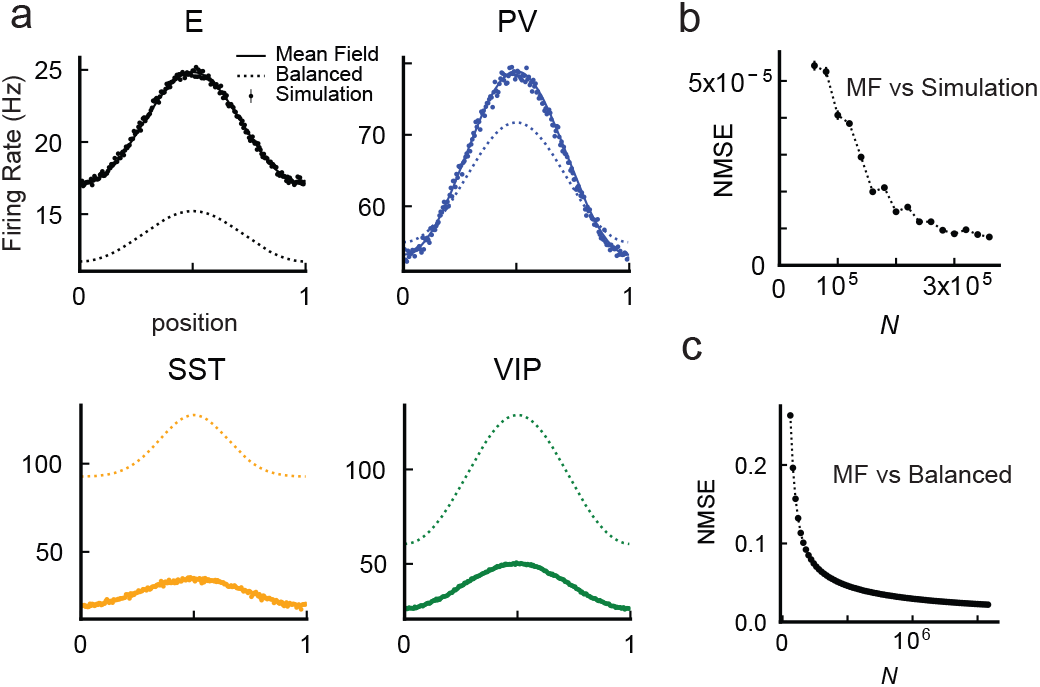
Firing rates in the four-population (E, PV, SST, and VIP) model. (a) Stationary firing rate from mean-field theory (solid lines), analytical solutions in the thermodynamic limit (dashed lines), and firing rates calculated from simulations (error bars) from spiking networks with *N* = 3 × 10^5^. For simulations, each data point represents the mean firing rate of neurons within a window of size 0.05; error bars correspond to mean ± sem over 10 independent network initializations. (b) Normalized mean square error (NMSE) between meanfield theory and simulated firing rates as a function of network size *N* . (c) NMSE between the mean-field model with varied network sizes and the balanced solution in the thermodynamic limit.

### D. Stability Analysis

One of the key features we include in the model is the long-range projections of SST neurons, i.e. *σ*_*s*_ *> σ*_*p*_, *σ*_*v*_. This is based on anatomical evidence as well as fact that long-range SST projections are necessary in spiking neural network models to reproduce the divisive effects of lateral inhibition by distal SST neurons [28]. However, previous work on spatially distributed E-I networks with a single type of inhibitory population showed that inhibitory projections must be strictly shorter than excitatory projections (*σ*_*i*_ *< σ*_*e*_) to maintain a stable balanced state [13, 14]. Here we examine whether long-range SST projections in a network composed of different inhibitory subtypes cause instability in our proposed model.

To do this, we linearize the firing rate dynamics about the fixed point,

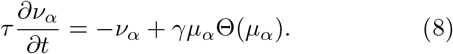

where *τ* is the characteristic time scale of the network, *γ* = 1 is the gain of the neuron, and Θ is the Heaviside step function. This approximates the network dynamics as threshold linear, which is accurate for high inputs [13, 35, 38, 39]. Eq. 8 thus describes the position-dependent linearized dynamics about the fixed point. We note that although the fixed point will depend on the static inputs 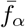 , its stability will not depend on 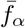 as long as the fixed point is strictly positive. Under this approximation, the fixed point is stable whenever the matrix 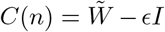 (where 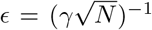 and *I* is the identity matrix) has eigenvalues *λ*_*n*_ with negative real parts Re(*λ*_*n*_) for each Fourier mode *n*.

We first consider the stability of a network of size *N* = 3 × 10^5^ with a homogeneous inhibitory neuron population by setting *q*_*i*_ ≡ *q*_*p*_ = 0.2 and *q*_*s*_ = *q*_*v*_ = 0. This is similar to the model in Ref. [13], but with inhibitory neuron parameters specifically derived from PV neurons. We computed the maximum real part of the eigenvalues for varying excitatory and inhibitory projection range *σ*_*e*_ and *σ*_*i*_ (Fig. 3a, top panel). The dashed line indicates the boundary between stable and unstable networks. When *σ*_*e*_ = 0.05 and *σ*_*i*_ = 0.2 (purple asterisk on Fig. 3a), the network lies in the unstable region. We then considered the stability diagram of the network with heterogeneous inhibitory neurons and vary *σ*_*s*_ along with *σ*_*e*_ while keeping *σ*_*p*_ = *σ*_*v*_ = 0.1 (Fig 3a, bottom panel). The boundary of stability has a significantly different profile. When *σ*_*s*_ = 0.2 and *σ*_*e*_ = 0.05 (black asterisk on Fig. 3a), stability is maintained.

**FIG. 3.**
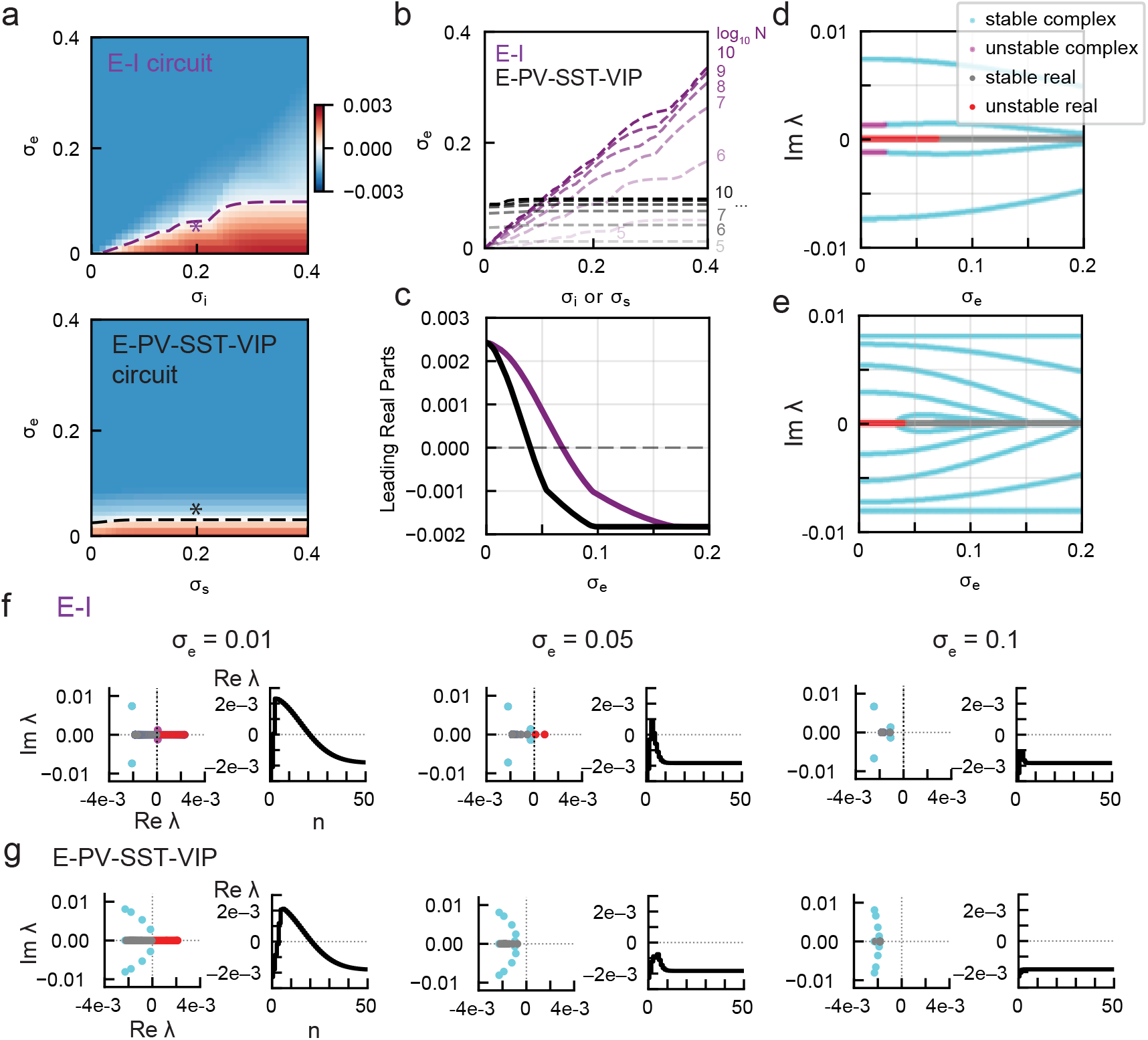
Stability in networks with homogeneous and heterogeneous inhibitory neuronal populations. (a) Stability diagrams of an E-I circuit with a homogeneous inhibitory neuron population consisting of PV neurons only (top) and of the E-PV-SST-VIP circuit with heterogenous inhibitory neurons (bottom; *N* = 3 × 10^5^). The color indicates the maximum value of Re(*λ*_*n*_) for the corresponding inhibitory projection distance *σ*_*i*_ and excitatory projection distance *σ*_*e*_. Red indicates positive eigenvalues (unstable) and blue negative values (stable). The dashed lines trace the borders of stability. (b) Stability boundaries for E-I (purple dashed lines) and E-PV-SST-VIP circuits (black dashed lines) for varying network sizes *N* (from 10^5^ to 10^10^, with darker shades indicating higher *N*). (c) Leading real eigenvalue part for varying *σ*_*e*_ for the two circuits. *σ*_*i*_ = 0.2 for the homogeneous circuit (purple) and *σ*_*s*_ = 0.2 for the heterogeneous circuit (black). (d) Imaginary parts of eigenvalues for varying *σ*_*e*_ in the homogeneous circuit. Pure real eigenvalues are colored gray (stable) and red (unstable) while complex eigenvalues are cyan (stable) and purple (unstable). (e) Same as (d) for the heterogeneous circuit. (f) Eigenspectrum *λ*_*n*_ for the homogeneous E-I circuit for *σ*_*e*_ = 0.01 (left panel), *σ*_*e*_ = 0.05 (center), and *σ*_*e*_ = 0.1 (right). In each panel, the left plot shows the eigenspectrum on the complex plane with the same color coding as in (d-e) and the right plot shows the real part of the eigenvalue for varying Fourier component *n*. (g) Same as (f) for the E-PV-SST-VIP circuit.

Increasing *N* for both networks increases the range of instability (Fig. 3b). For the homogeneous model, the stability-instability boundary trends towards the diagonal line *σ*_*e*_ = *σ*_*i*_. This indicates that as *N* → ∞, the network is unstable as long as *σ*_*i*_ *> σ*_*e*_. However, for the heterogeneous network the boundary converges to a horizontal line, and a region exists for which the network remains stable when *σ*_*s*_ *> σ*_*e*_. Network stability is thus insensitive to the broadness of SST projections. In contrast, the broadness of PV projections in the heterogeneous network causes instability even when SST and VIP neurons are kept short-range (See SFig. 1 [40]), indicating that projections from PV neurons specifically are crucial in maintaining dynamical stability of the network. Cell-type-specific connectivity underlies this effect, and in particular PV self-inhibition is necessary for its stabilizing role, as demonstrated by a decomposition analysis in the Supplemental Material, Sec. 1 [40].

The E-PV-SST-VIP circuit hence has a larger region of dynamical stability. This is seen explicitly in Fig. 4c, which displays the real parts of the eigenvalue when inhibitory projections are fixed and *σ*_*e*_ varied. Instability occurs when *σ*_*e*_ is below 0.07 for the homogeneous model and below 0.04 for the heterogeneous model (Fig. 3c). The two models show distinct imaginary components, as seen in Figs. 3d, e. Furthermore, in both the homogeneous and heterogeneous model, the first emergent unstable eigenvalues are pure real (red line). The homogeneous model additionally displays unstable complex eigenvalues at very narrow *σ*_*e*_.

**FIG. 4.**
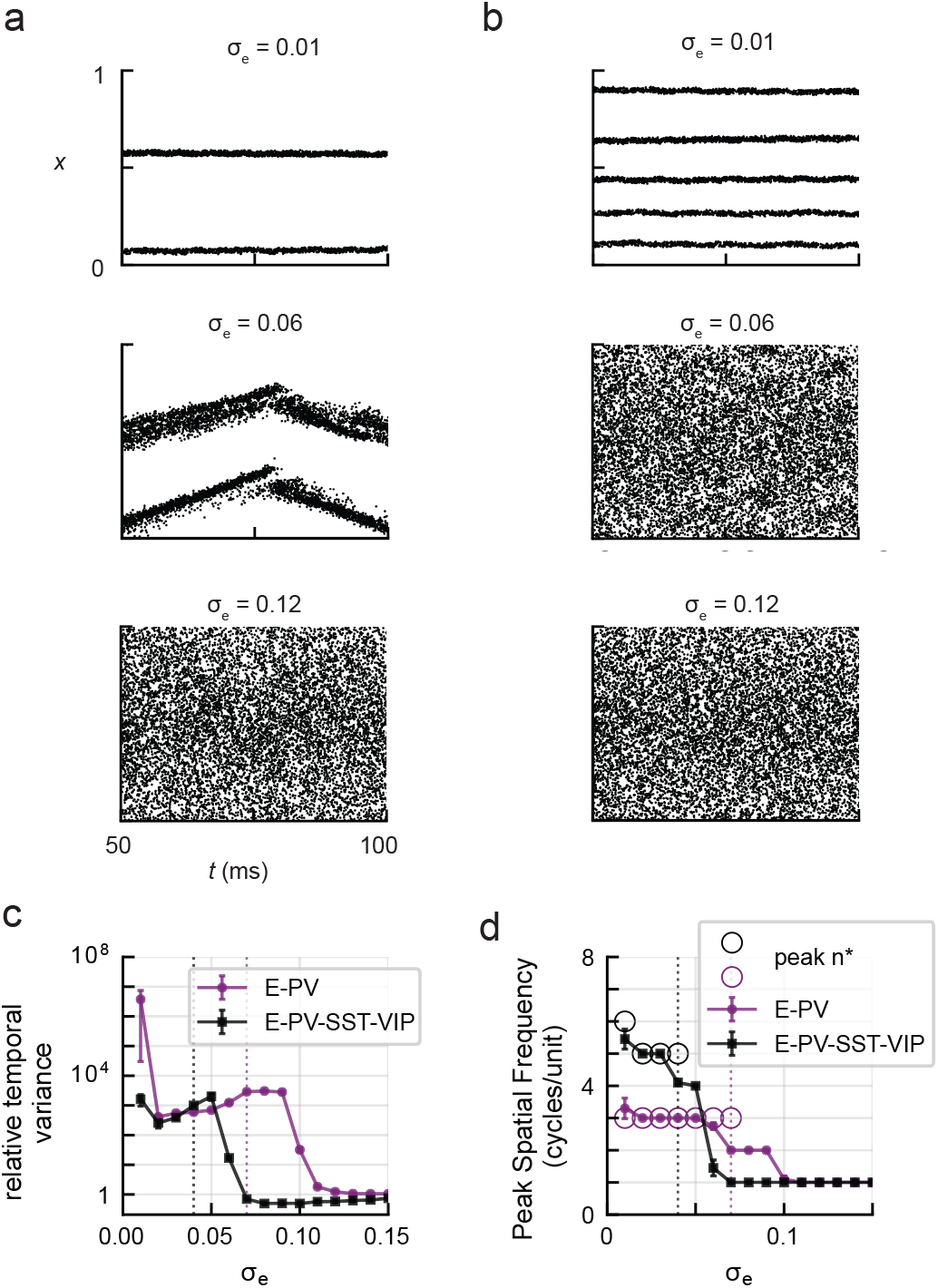
Spatiotemporal network activity in spiking network simulations. (a) Raster plot of excitatory neurons in the E-I circuit for *σ*_*e*_ = 0.01 (top), 0.06 (middle), and 0.12 (bottom) in response to Gaussian input. For visibility, only spikes from one in every eight neurons is shown. (b) Same as (a) for the E-PV-SST-VIP circuit. (c) Mean relative temporal variance of firing rates red of a sample of 100 E neurons in the two circuits, for varying *σ*_*e*_. Raw variance is normalized against the variance when *σ*_*e*_ = 0.2. Shown are the mean ± sem over 20 independent simulations. Vertical dotted lines show the bifurcation points from Fig. 3a. (d) Peak spatial frequency from simulations (purple dots for E-I, black dots for E-PV-SST-VIP) of firing rate activity in the two models for varying *σ*_*e*_. Open circles indicate the peak unstable frequency *n*^*∗*^ from mean-field theory. Across all panels, *σ*_*i*_ = 0.2 for the E-I model and *σ*_*s*_ = 0.2 for the E-PV-SST-VIP model.

To further characterize the bifurcation, eigenspectra of *C*(*n*) are shown for three values of *σ*_*e*_ in Fig. 3f (E-I circuit) and Fig. 3g (E-PV-SST-VIP circuit). Each pair of panels show the eigenspectrum on the complex plane on the left and the real eigenvalue parts for varying Fourier node *n*. In both models, the critical Fourier node *n*^*c*^ that initially becomes unstable (Re *λ*(*n*^*c*^) ≥ 0) is strictly positive, indicating a Turing bifurcation.

### E. Spatiotemporal features of network dynamics

We then proceeded to verify the spectral analysis of the mean-field model in Fig. 3 with spiking neural network simulations. Spatially-organized raster plots, shown in Fig. 4a for the E-I circuit, show the emergence of spatial patterns for *σ*_*e*_ = 0.01 and 0.06; when *σ*_*e*_ = 0.12, the firing pattern reflects the smooth Gaussian input. For the E-PV-SST-VIP circuit in Fig. 4b, we also see spatial patterns when *σ*_*e*_ = 0.01 but with a higher spatial frequency, which disappear when *σ*_*e*_ = 0.06.

We quantified the temporal variance of firing rates of the two models for varying *σ*_*e*_ in Fig. 4c. Specifically, we quantified the temporal variance in the firing rates of neurons within local patches of the network (see Sec. IV E for details). As predicted by the transition points in Fig. 3c, temporal variance increases rapidly as *σ*_*e*_ approaches the critical *σ*_*e*_ predicted by the stability analysis (dotted vertical lines, Fig. 4c), which is higher for the E-I circuit compared to the E-PV-SST-VIP circuit. We note that for both models, the region of high temporal variance occurs at a higher *σ*_*e*_ than predicted by the theory, which may be due to finite size effects.

We further examined the emergent spatial patterns in the models. Fig. 4d displays the peak spatial frequency in the two models given by the peak frequency of firing rate activity over a 20 ms window of activity (Sec. IV E). When *σ*_*e*_ is well above the bifurcation point (*>* 0.06 for the heterogeneous and *>* 0.09 for the homogeneous circuit), the networks track the spatial frequency of the Gaussian input with a frequency of 1 cycle per spatial unit. Spatial patterns emerge in both models as *σ*_*e*_ is decreased. In the unstable region, we expect the Fourier nodes *n* with unstable eigenvalues to strongly influence the spatial frequency of the resulting dynamics. We thus compared the spatial patterns in simulations against the peak unstable Fourier modes *n*^∗^ (peaks of black lines in Figs. 3f, g) represented by open circles in Fig. 4d. In each model, for *σ*_*e*_ smaller than the critical point, the peak spatial frequencies observed in simulations closely match the predicted *n*^∗^. When close to the bifurcation points, there is a competition between the frequency of the Gaussian input and the dominant Fourier mode, and the peak spatial frequency has an intermediate value.

### F. Cell-type-specific gain modulations

Next, we examined the gain modulations induced by stimulating specific cell types in the mean-field model. We define gain modulations as changes in the slope of the visual response curves in excitatory neurons: a decrease in the slope indicates a divisive modulation, while a vertical shift without slope change indicates a purely subtractive modulation [22, 26, 28]. We presented a Gaussian-shaped input current to excitatory and PV neurons centered at *x*_*v*_ = 0.5 with varying intensities to simulate visual inputs of different contrast levels. Cell-type-specific modulatory signals activating inhibitory neurons were induced with an additional Gaussian-shaped input current centered at *x*_*C*_ which was either proximal (*x*_*C*_ = 0.5) or distal (*x*_*C*_ = 0) to the site of the visual stimulus input (Fig. 5a; details in Sec. IV F). The spatial structure of the E response with PV (blue), SST (gold) and no stimulation (black) showed distinct shapes with proximal stimulations (Fig. 5b, left). However, the slope of excitatory responses to varying stimulus intensity did not decrease when paired with stimulation of any types of inhibitory neurons, i.e. no divisive gain changes were observed (Fig. 5b, right). Different spatial response profiles were also observed with distal stimulation of inhibitory subtypes, but again, with no divisive modulations (Fig. 5c).

**FIG. 5.**
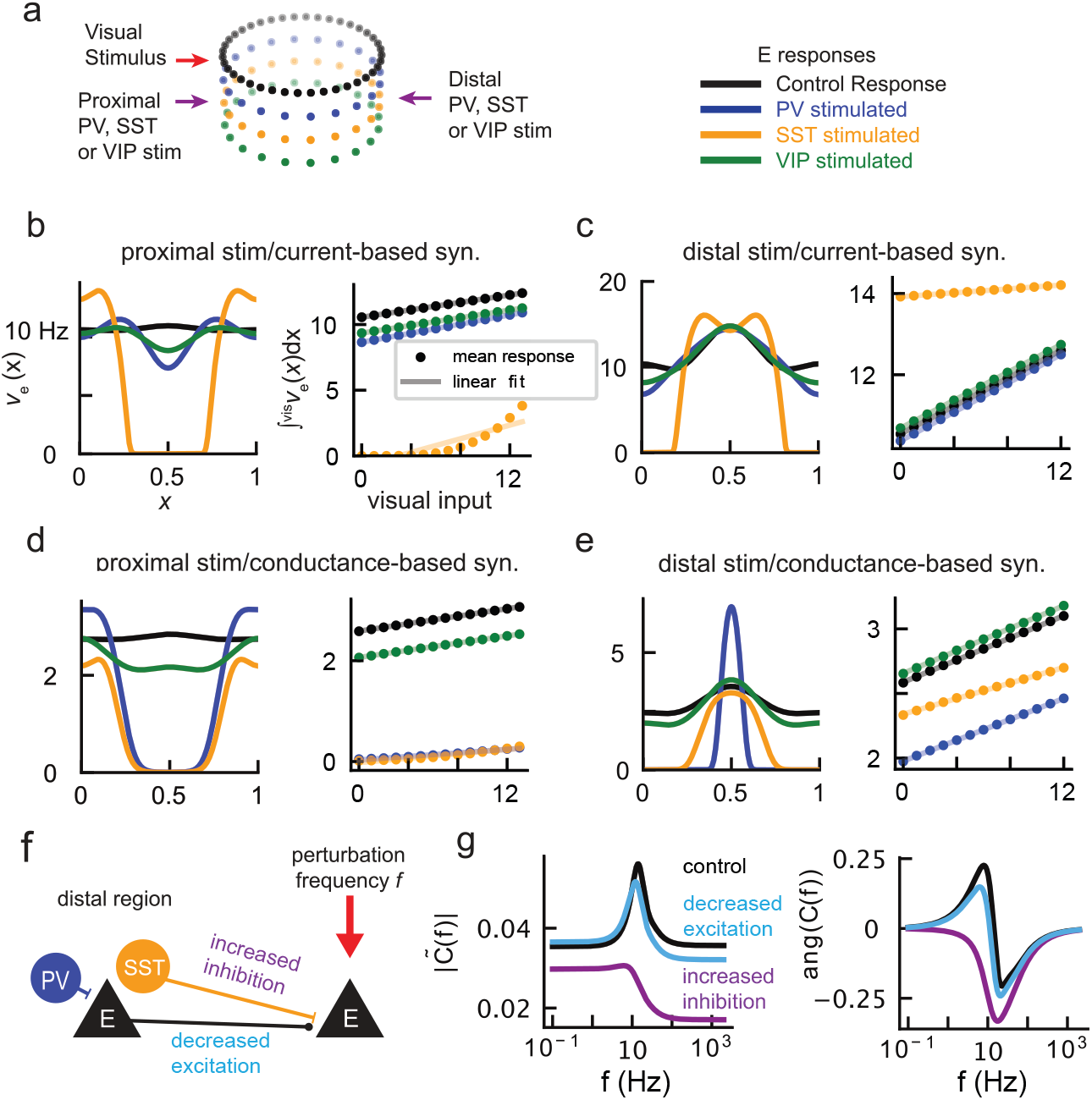
Cell-type-specific gain modulations. (a) Schematic of stimulations. Visual stimulus is given as input to E and PV neurons centered at *x*_*v*_ = 0.5. For proximal modulations, PV, SST, or VIP neurons at *x*_*v*_ = 0.5 were stimulated. For distal modulations, neurons at *x*_*C*_ = 0 were stimulated. (b) Left: Response profiles of E neurons in the current-based model with PV (blue), SST (gold), VIP (green) proximal stimulation and without activation of inhibitory neurons (black). Right: Dots represent E responses averaged across the region of visual input for varying input strength (see Eq. 29). Shaded lines show linear fits. (c) Same as (b) but for distal PV, SST, and VIP stimulation. (d, e) Same as (b, c) but with the conductance-based model. (f) Schematic of linear response analysis. Oscillatory perturbation is applied to the visual input. The linear response with increased long-range inhibition (purple) and decreased long-range excitation (cyan) are computed and compared against the unaltered network conditions (black). (g) Linear response amplitude (left) and phase shift (right).

Experimental evidence suggests that both PV and SST neurons can exert divisive gain modulations (i.e., negative slope changes) depending on the stimulation conditions and the location of their stimulation relative to the site of visual input [23–25, 28]. We reasoned that the lack of divisive gain modulations in the model may be due to simplifications in the model, in particular due to the current-based synapses assumed in the model. This is manifested in the balanced solution Eq. 5, which is linear in external inputs 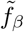. We thus incorporated conductance-based synapses into the model; mean-field firing rates can be computed with a similar approach to the current-based case with modified parameters and a voltage-dependent diffusion term (see Sec. IV D). We first verified that the mean-field results with the conductance-based synapses agreed with the corresponding spiking network simulations (SFig. 2 [40]). We repeated the stability analysis across spatial scales of excitatory and inhibitory projections to account for voltage-dependent synaptic transmissions and found similar results to the current-based model (SFig. 3 [40]).

Having verified that the conductance-based meanfield model generates the same results on spatial response profiles and projection-dependent stability as with the current-based synapses model, we next simulated proximal and distal activations of inhibitory neurons using the mean-field model with conductance-based synapses. When proximal stimulation was given to the conductance-based network, again distinct profiles emerged—PV and SST stimulation completely suppressed excitatory responses at the site of visual input, while VIP stimulation exerted a subtractive effect (Fig. 5d). With distal stimulation, the spatial profile of the excitatory response sharpened with PV stimulation. In addition, distal PV stimulation caused a subtractive effect on the E response to visual input strengths (no slope change), while distal SST stimulation caused a divisive gain change (decrease in slope). Distal VIP stimulation caused slight multiplicative gain change (increase in slope, Fig. 5e). These distance- and cell-type dependent modulations are consistent with experimentally observed effects of both proximal and distal PV or SST stimulation in mouse V1: both resulted in strong divisive gain modulations in excitatory neurons when the activation of the inhibitory population was proximal, and only SST activation resulted in divisive operations when distal [28]. The positive gain change induced by VIP neurons is also consistent with a previous experimental study [31].

### G. Direct inhibition promotes divisive gain modulations

Inhibitory neurons can exert gain modulations through multiple pathways [21]. Here we examine whether the direct inhibition or indirect disexcitation may underlie the distinct modulatory effects. Similar to the network response, the slope of an input-output curve of a single neuron can be interpreted as neuronal gain [22]. It is known that the background synaptic input affects the gain of a neuron’s response [41]. We can thus trace the gain modulation of the network to shifts in the background inputs of neurons due to cell-type-specific activations. Distal SST neurons, due to their long-range projections, can directly inhibit E neurons in the visual field of stimuli. On the other hand, distal PV neurons primarily act on E neurons in the visual field polysynaptically by inhibiting presynaptic E neurons, thereby decreasing excitation (Fig. 5f). Within the mean-field framework, linear response theory can be used to measure the precise response of neurons to perturbations in the input, giving us a precise way to measure neuronal gain.

We used linear response theory to quantify the neuronal gain of excitatory neurons in the network at the center of the visual input. Specifically, we examined how the gain changed when direct inhibition was increased (simulating the distal SST effects) versus when the background excitatory input was decreased (simulating the distal PV effects). A frequency-dependent perturbation was introduced to the visual input in the two cases as well as to the unmodified control (details in Sec. IV G). Inhibition was increased and excitation decreased each up to 10-fold, with results summarized in SFig. 4[40]. Here we examined the amplitude and angle of the response with a 2-fold increased inhibition and 10-fold decreased excitation, and compared them to the base case (Fig. 5g). The amplitude of the response in the control case showed a peak at 10 Hz, indicating a resonant frequency, and there was an associated positive phase shift (black curve, Fig. 5g). When excitation was decreased, the response was similar to the control case, exhibiting the same resonant frequency and phase shift (cyan curve). However, when inhibition was increased, the amplitude of the response was shifted downward, indicating a lower gain (purple curve). Moreover, the resonant frequency disappeared with increased inhibition. These results indicate that increased inhibition induces a negative gain change in the excitatory neurons across all stimulus frequencies, which may act as a mechanism for divisive effects of distal SST stimulation. Distal PV stimulation decreases excitation without increasing direct inhibition, resulting in a subtractive effect. Meanwhile, a strong local stimulation of either PV or SST can induce divisive operations through direct inhibition.

## III. DISCUSSION

We derived mean-field equations for a spatially-organized spiking neural network with excitatory and three inhibitory sub-populations. The neurons have distance-dependent connectivity rules that depend on pre- and post-synaptic cell types, informed by anatomical data from mouse primary visual cortex. Given the cell-type specific connectivity, we derived conditions for E-I balance in the network. The model revealed dynamical properties conferred by inhibitory heterogeneity that may indicate computational benefits. The heterogeneous network maintained stable dynamics even with long-range SST inhibition; this was in contrast to the network with homogeneous inhibitory neurons, whose projections were restricted to be shorter than those of excitatory neurons in order to maintain stability as suggested by a previous theoretical study [13]. Thus, our result, by introducing inhibitory subtypes, closes the gap between the previous theoretical prediction where inhibitory projections are restricted to be local [13] and contradictory experimental observations of long-range inhibition [28– 30]. Furthermore, activation of each inhibitory cell type performed distinct modulations on excitatory neurons processing visual input, in agreement with recent experimental observations [25, 28, 31]. This demonstrates that there are multiple modes of computations that can be activated flexibly depending on spatial relation between inhibitory modulatory signals and visual stimuli.

Our work sheds light on the computational role of neuronal cell type diversity and their distinct spatial connectivity in the cortex. The balance of excitation and inhibition in the cortex is believed to be the basis for irregular spiking activity and heterogeneity of responses [8]. By extending previous work on spatially-distributed E-I networks [13, 15], we demonstrated that the network composed of excitatory and three main subtypes of inhibitory neurons—PV, SST, and VIP—can achieve balance to each of its populations. Our analytical findings give us a simple description to determine balance in a multi-population network depending on connectivity and synaptic weights. Understanding how balanced states can be altered by changes in connectivity may be important in understanding network activity in neurological disorders [42–44].

Our mean-field approach further allowed us to directly compute the eigenspectrum of the network’s fixed state. Our model shows that neural heterogeneity enhances network stability, allowing stable firing rate readouts even in the presence of long-range inhibition. In contrast, networks with a homogeneous inhibitory population displayed unstable dynamics and high response variability with long-range inhibition. This result thus implicates possible computational advantages of cell type diversity. Loss of stability was caused by Turing bifurcations in both heterogeneous and homogeneous models. Simulations confirmed the emergence of spatial patterns as excitatory projections became local, with a higher spatial frequency exhibited in the heterogeneous model.

Identifying neuronal cell types and their functional roles has emerged as a major problem in neuroscience [17, 45, 46]. Computational studies have played an indispensable role in understanding the role of inhibitory neuron subtypes in dynamics and computations [18–20, 32, 47– 49]. Our work complements these studies by providing additional mechanisms by which networks may perform divisive and subtractive computations. For example, Wilmes et al. [20] showed that divisive and subtractive operations on excitatory rates by PV and SST neurons, respectively, may account for computations needed to encode the variance and mean of a variable. Recent theoretical work by Bos et al. analyzed the multiple pathways through which SST modulate E neurons and showed that SST activation can exert differential gain modulations on excitatory neurons depending on the synaptic weights [21]. Our work complements these studies by showing that PV and SST neurons can perform various degrees of divisive or subtractive modulations depending on the distance between their activation and the stimulus input sites. Thus, inhibitory neurons, by utilizing their heterogeneous projection patterns across the cortical network, may perform a range of distinct operations on spatially organized stimuli in an effective and robust manner. The linear response analysis suggest that a shift in the balance of excitation and inhibition may underlie divisive gain changes, with direct inhibition contributing to divisiveness.

Our results highlight the computational benefits of a heterogeneous neuronal population in recurrent neural networks. A recent line of work has found that the degree of heterogeneity within a cell type can play a major role in network computations [48, 49]. While our work focused on heterogeneity in intrinsic properties and projection patterns across cell types and assumed no variation in parameters within each cell type, it would be interesting to investigate the interaction of within- and between-population heterogeneity. In another work, Winston et al. found that heterogeneous cell types emerge when training spiking neural networks to perform temporal processing tasks [47]. It is an open question whether PV, SST, and VIP-like structures may emerge from network optimization under certain objective functions and constraints.

The current work can be expanded in several ways. One line of study would be to investigate how the network can take computational advantage of the response properties explored here by employing them in visual processing tasks. In addition, further biophysical details neglected in the present model, such as dendritic integration [28], can be incorporated to make the mean-field model more realistic.

## IV. METHODS

### A. Neuronal dynamics and spatial projections

The membrane potential *V*_*α,k*_ of the *k*th neuron of type *α* (= *e, p, s*, or *v*) is given by leaky integrate-and-fire (LIF) equations

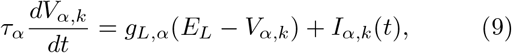

where *τ*_*α*_ is the membrane time constant, set to 30 ms for *α* = *e* and 20 ms otherwise; *g*_*L,α*_ is the leak conductance, set uniformly to 1 (dimensionless units); *E*_*L*_ is the leak reversal potential at 0; and *I*_*α,k*_(*t*) is the input current, given by

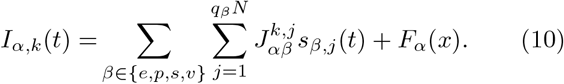

Here 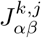 is the synaptic weight from presynaptic neuron *j* of type *β*, 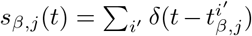 is its spike train, *x* = *k/N*_*α*_ is the position of the *k*th neuron, and *F*_*α*_(*x*) is a static, position-dependent external input. The membrane potential variable *V* and voltage-related parameters are given in dimensionless units.

When the membrane potential reaches the threshold *V*_th_, a spike is emitted and *V*_*α,k*_ is reset to the reset voltage *V*_re_ on the next time step. We set *V*_th_ = 1 and *V*_re_ = 0; numerical integration is performed using the Euler method with time step Δ*t* = 0.1 ms

The synaptic weight 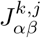 from neuron *j* of cell type *β* (at position *y*) to neuron *k* of cell type *α* (at position *x*) is given by

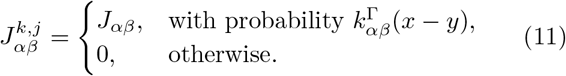

Here 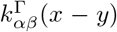 is the distance-dependent connectivity kernel.

The synaptic weight *J*_*αβ*_ is assumed to scale with the network size as 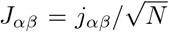 where *j*_*αβ*_ is a constant independent of *N* . External inputs scale according to 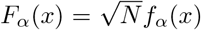, while other parameters are constant with respect to network size.

We assume that the connectivity kernel is a wrapped Gaussian projection kernel with a broadness that depends on the presynaptic neuron type, i.e., 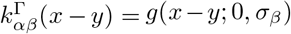 , where the wrapped Gaussian *g* is given by

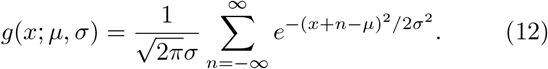

Gaussian spatial connectivity profiles have been used to fit anatomical data of rodent and cat cortex [17, 50, 51]. The Gaussian model breaks down for long-range connections beyond approximately 1 mm [52] which is beyond the range considered in the model. The assumption that the spatial connectivity kernel only depends on the cell type of the presynaptic neuron is a simplification based on evidence that the primary constraint on distance-dependent connectivity is the axonal projections of the presynaptic neurons [53, 54]. We note however that this assumption is not strictly necessary. Current inputs are given by

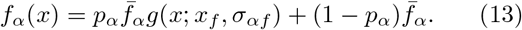

Here 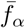 is the mean input to cells of type *α* and *p*_*α*_ determines the proportion of the input spatially localized at *x*_*f*_ .

### B. Mean-field equations in the continuum limit

In the mean-field model, we take the continuum limit

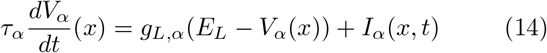

where the input current from Eq. 9 is replaced by a position-dependent effective field, modeled as a random process 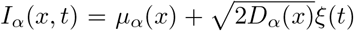. Here *µ*_*α*_(*x*) is the mean input current, given by a sum of recurrent inputs and position-dependent current input *f*_*α*_ (see Eq. 2),

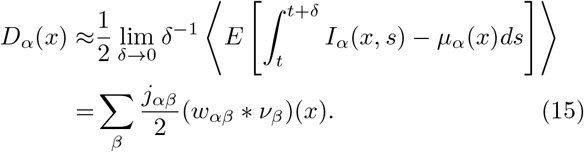

is the input variance, and *ξ*(*t*) is zero-mean delta-correlated white noise. Here 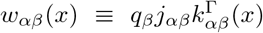 is the distance (*x*)-dependent weighted connectivity, *ν*_*α*_(*x*) ≡ ⟨*E*[*s*_*α,k*_]⟩ is the population- and position (*x*)-dependent mean firing rate, and ∗ indicates circular convolution, i.e. (*w*_*αβ*_ ∗ *ν*_*β*_)(*x*) ≡ ∫_Γ_ *w*_*αβ*_(*x* − *x*′)*v*_*β*_(*x*)*dx*′.

Given input mean and variance, the firing rate of neurons of type *α* in position *x* is computed using a Fokker-Planck approach [36, 37]. For a given *α* and *x*, the variables *p*_*s,α*_(*x, V*) = *P*_*s,α*_(*x, V*)*/ν*_*s,α*_(*x*) and *j*_*s,α*_(*x, V*) = *J*_*s,α*_(*x, V*)*/ν*_*s,α*_(*x*), where *P*_*s,α*_(*x, V*), *J*_*s,α*_(*x, V*), and *ν*_*s,α*_(*x*) are the stationary probability distribution of the membrane potential, the current operator, and the firing rate, respectively, are described by

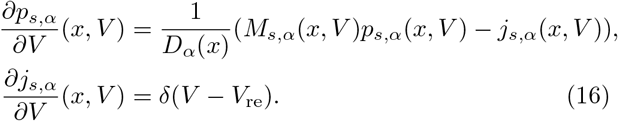

Here 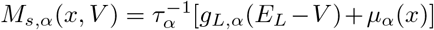 is the drift coefficient. We integrate Eqs. 16 using a numerical integration scheme previously introduced [36], specifically using Simpson’s rule [37]. The firing rate is then given by

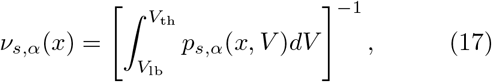

where the membrane potential lower bound *V*_lb_ is selected so that *P*_*s,α*_(*x, V*_lb_) is negligible (here set to *V*_lb_ = −1).

This numerical integration scheme defines a mapping Φ from the input mean *µ*_*α*_(*x*) and variance *D*_*α*_(*x*) to the position-dependent firing rate, defining a set of self-consistent equations *ν*_*α*_(*x*) = Φ(*µ*_*α*_(*x*), *D*_*α*_(*x*)). Fixed points are found using an iterative scheme as described in Ref. [37]. Starting from an initial guess 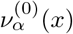 for each *α*, 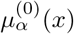 and 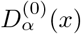 are computed using Eq. 2 and 15 and the next guess is computed iteratively,

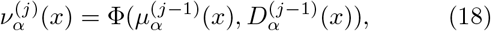

until the L2 deviation between 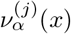 and 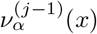 is less than a threshold, which we set to 10^−7^ Hz.

### C. Balanced state in the thermodynamic limit

Theories of randomly connected neural networks sought to explain asynchronous spiking activity with models of a balanced state in which excitatory and inhibitory inputs dynamically achieve balance [35]. In this state, firing rates of neurons can be expressed in analytic form which becomes exact in the large-*N* limit. In Sec. II B we generalize the approach by Rosenbaum and Doiron [13] which implemented two populations—one excitatory and one inhibitory—for the four-population model where the inhibitory population is split into three subtypes. Here we provide details about derivation of Eq. 6 in Sec. II B.

Each component of 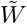 in Eq. 5 can be written 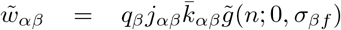 where *q*_*β*_, *j*_*αβ*_, 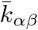 and are constants independent of *n* and 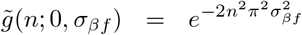 is the *n*th Fourier component of the wrapped Gaussian. We can thus write 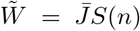 where 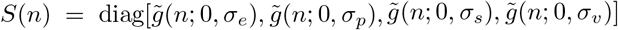 and 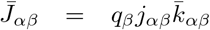, yielding 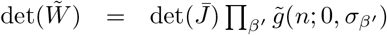. The cofactors have a similar structure and can be written 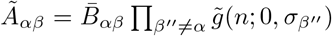, where 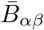 are cofactors of 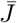. Hence,

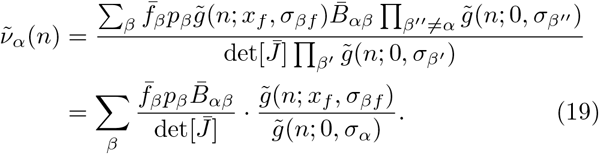

We note that because of the cancellation of factors 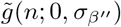 in Eq. 19, the spatial profile of population *α* will not depend on *σ*_*β*_ for *β* ≠ *α*, provided that the solution exists. In order to have lim_*n*→∞_ 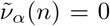, we must have *σ*_*βf*_ *> σ*_*α*_ for each *α, β*. If this is satisfied, the balanced solution can be written as

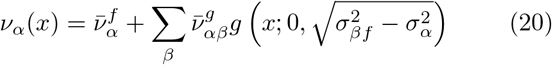

where

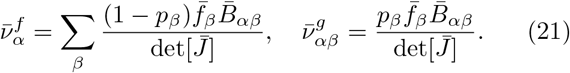

For this to be a valid physical solution, the firing rates in Eq. 20 must be positive. This does not imply that each term in Eqs. 21 must be positive. The generalized solution admits solutions where some terms are negative, resulting in inverted (but positive) firing rate profiles.

Synaptic weights *j*_*αβ*_ and external input intensities 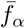 that achieve balance are generated numerically as follows. Synaptic weights are constrained to be in the range [0.2, 2] and external inputs in the range [10^−5^, 10^−1^]. Initial values are generated randomly within the bounds and then the parameters are adjusted to minimize the mean square deviation between 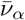 and target firing rates set at 50 Hz using the Sequential Least Squares Programming algorithm. If the resulting firing rates are all positive, the parameters are accepted. One such set of synaptic weights and external input parameters was used Figs. 1-4 and are listed in Table II. Parameters used in Fig. 5 are described in Sec. IV F.

### D. Conductance-based model

When simulating conductance-based neurons, Eq. 10 becomes

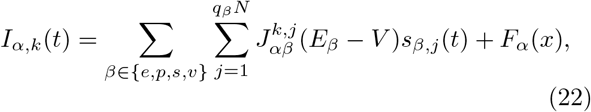

where *E*_*β*_ is the reversal potential, set to *E*_*e*_ = 1.5 for excitatory and *E*_*β*_ = *E*_*i*_ = −0.5 for inhibitory presynaptic neurons. For conductance-based equations, we replace *E*_*L*_ in Eq. 16 with the effective leak voltage 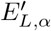 which is dependent on cell type [36],

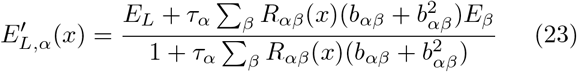

and the membrane conductance *g*_*L,α*_ with

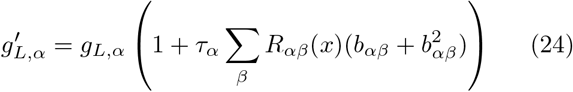

where 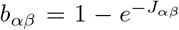 and *R*_*αβ*_ is the input spike rate from population *β*, given by

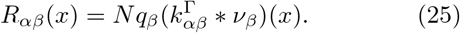

We use the same neuronal and synaptic parameters *g*_*L,α*_, *τ*_*α*_, and *j*_*αβ*_ as in the current-based case (Table II). We integrate Eqs. 16 as before, now with a voltage-dependent diffusion

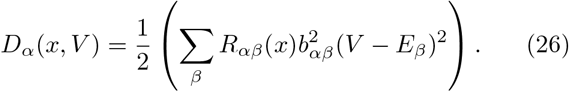

Due to inputs being voltage-dependent, the stability matrix 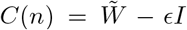, where 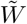 is the matrix with Fourier components 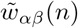, is also modified. We make the simplifying assumption that the mean membrane potential is independent of *x*, yielding

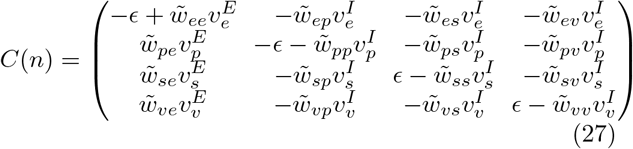

Where 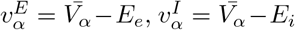, and 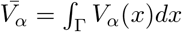.

### E. Spatiotemporal analysis

Temporal variance was computed to quantify the variability in firing-rate readouts over time. For each simulation, we selected five equally-spaced patches of size 0.1 (i.e., (0, 0.1), (0.2, 0.3), etc.). For each patch, the spike times of all E neurons in the patch were binned into 5 ms temporal windows over the analysis period (100 ms with the initial 100 ms excluded to remove transient effects) and converted to instantaneous mean firing rates. The variance of the firing rate across time was computed for each patch and the variances were averaged across the patches. This procedure was repeated for 20 random seeds. For each *σ*_*e*_ value, the mean temporal variance was calculated by averaging across all seeds.

To estimate the peak spatial frequency of population activity, we first formed a spatial firing-rate profile by binning spikes that occurred within a 20 ms window across the position coordinate *x* ∈ [0, 1] with bin width Δ*x* = 0.02 and computing the firing rate within each bin. The mean was subtracted from the resulting spatial rate profile and its discrete spatial Fourier transform was computed via a Fast Fourier Transform (FFT). The one-sided spatial power spectrum was then formed by retaining only positive spatial frequencies (in cycles per unit distance). The peak spatial frequency was defined as the frequency with maximum power. This process was again repeated for the 20 random seeds.

### F. Modulation of visual response

Modulations by specific inhibitory neuronal populations are measured by changes in the excitatory neurons’ response to a visual stimulus. We first set *p*_*α*_ = 0 in Eq. 13 to induce spatially uniform input 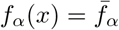 and the network parameters 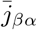 and 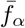 are re-tuned to bring the network firing rate to below 10 Hz, which are the observed spontaneous firing rates in mouse V1 in the absence of visual stimuli [28]. The resulting parameters are listed in the Supplemental Material, Sec. 2 [40]. Visual stimuli and modulatory signals are given in addition to the baseline input.

The visual input is incorporated by an additional term *I*_v,*α*_(*x*) to the total input given to the mean input in Eqs. 10 or 22. For the current-based model, this is given by

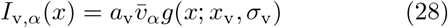

where *a*_v_ controls the overall strength of the visual input, 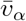 is a population-dependent term controlling the relative strength of the visual input to population *α, x*_v_ = 0.5 is the center of the stimulus and *σ*_v_ = 0.3 is the spatial spread of the input. Feedforward visual inputs are known to be fed to E and PV neurons [26], so we set 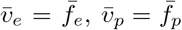, and 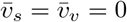.

For the conductance-based model, the visual stimuli are represented by conductance-based synaptic inputs. This results in an additional term in each of the summations over *β* in Eqs. 23, 24, 26 to include inputs from an external layer *T* where we set *R*_*αT*_ (*x*) = *a*_v_*r*_*α*_*g*(*x*; *x*_v_, *σ*_v_) for *α* ∈ *e, p* and *R*_*αT*_ = 0 otherwise. We set *r*_*e*_ = 4 × 10^−4^ and *r*_*p*_ = 1 × 10^−4^, values tuned to achieve realistic firing rate responses [28]. Synaptic weight parameters are set to *b*_*eT*_ = 1 − *e*^−0.2^ and *b*_*pT*_ = 1− *e*^−0.02^.

To measure modulations, we first measure the excitatory response of the system to visual stimuli of varying intensity without additional activation of inhibitory neurons. This is done by computing the cell-type and position-dependent stationary firing rates *ν*_*α*_(*x*) for *a*_v_ varying from 0 to 12. Then the process is repeated with an additional current input given to a specific inhibitory cell type, similar to optogenetic stimulations in experimental studies [25, 28, 31]. The inhibition-activating stimulus is given by a Gaussian current input centered at *x*_*C*_, which is set to *x*_*C*_ = 0.5 for proximal stimulations (Figs. 5b,d) and 0 for distal stimulations (Figs. 5c,e). The spatial spread of the inhibition-activating input is given by the standard deviation *σ*_*C*_ = 0.2 and the magnitude is set equal to 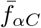, i.e., 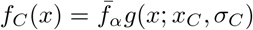. In the current-based model, 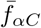 was first set to 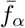 and then increased until the modulation partially silenced at least one population in order to induce nonlinear effects– otherwise, no gain modulations were observed. In the conductance-based model, the effects of modulation were stronger and the modulation was decreased if complete suppression of responses occurred. Parameters are listed in the Supplemental Materials, Sec. 2. The mean visual response (dots in Figs. 5d-e) is defined as the mean excitatory firing rate in the range [*x*_v_ − *σ*_v_, *x*_v_ + *σ*_v_], i.e.,

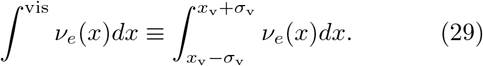

### G. Linear response theory

Linear responses to input perturbations are computed with the methods developed in Refs. [36, 37] and briefly outlined here. Linear responses are measured for excitatory neurons at position *x* = 0.5; cell type indices and positions are suppressed in the following equations. If a weak noise term 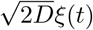 is added to the visual input *I*_v_(*t*) (we set *D* = 0.1), the cross-spectral density is given by

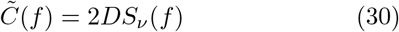

where *S*_*ν*_ (*f*) is the susceptibility function of the firing rate. We consider a perturbation to the visual input

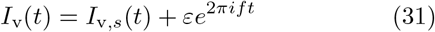

where *I*_v,*s*_(*t*) is the unperturbed input to the neuron, *f* is the frequency of the modulation and *ε* is a small parameter controlling the amplitude of the modulation. Setting *ε* = 0 recovers the fixed-point solution. To first order in *ε*, the response of the neuron is periodic with frequency *f* ,

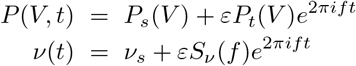

where *P*_*s*_(*V*) and *ν*_*s*_ are the steady-state probability density function and the firing rate, respectively, and *P*_*t*_(*V*) and *S*_*ν*_ (*f*) are the susceptibility functions. From here a time-dependent Fokker-Plank equation for the probability density *P* (*V, t*) can be derived with a drift coefficient

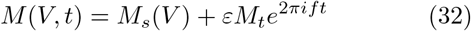

where *M*_*s*_(*V*) is the stationary drift coefficient in Eq. 16 and *M*_*t*_ is the coefficient of the time-dependent drift, given by *M*_*t*_ = *g*_*L,α*_*τ*_*α*_. We can decompose the probability density and flux functions as *P*_*t*_(*V*) = *S*_*ν*_ *p*_*ν*_ (*V*)+*p*_*t*_(*V*) and *J*_*t*_(*V*) = *S*_*ν*_ *j*_*ν*_ (*V*) + *j*_*t*_(*V*) which satisfy the equations

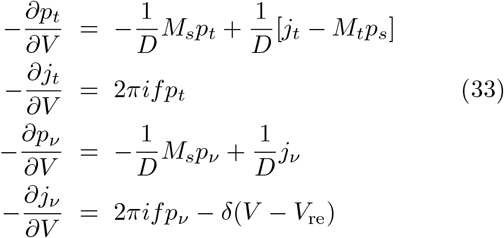

with boundary conditions *p*_*ν*_ (*V*_th_) = *p*_*t*_(*V*_th_) = *j*_*t*_(*V*_th_) = 0 and *j*_*ν*_ (*V*_th_) = 1. We integrate these equations with Simpson’s rule to obtain the susceptibility,

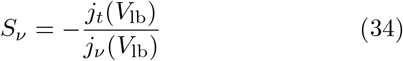

where *V*_lb_ is the lower bound of the membrane potential, set to −1. The cross-spectral density is then computed using Eq. 30.

## Data, Materials, and Software Availability

All code used in this study is available on https://github.com/HChoiLab/CellTypeMeanField.

## ACKNOWLEDGMENTS

This work was supported by the National Eye Institute of the National Institutes of Health under Award Number R00 EY030840. The content is solely the responsibility of the authors and does not necessarily represent the official views of the National Institutes of Health.

## Supplemental Material

**Figure S1:**
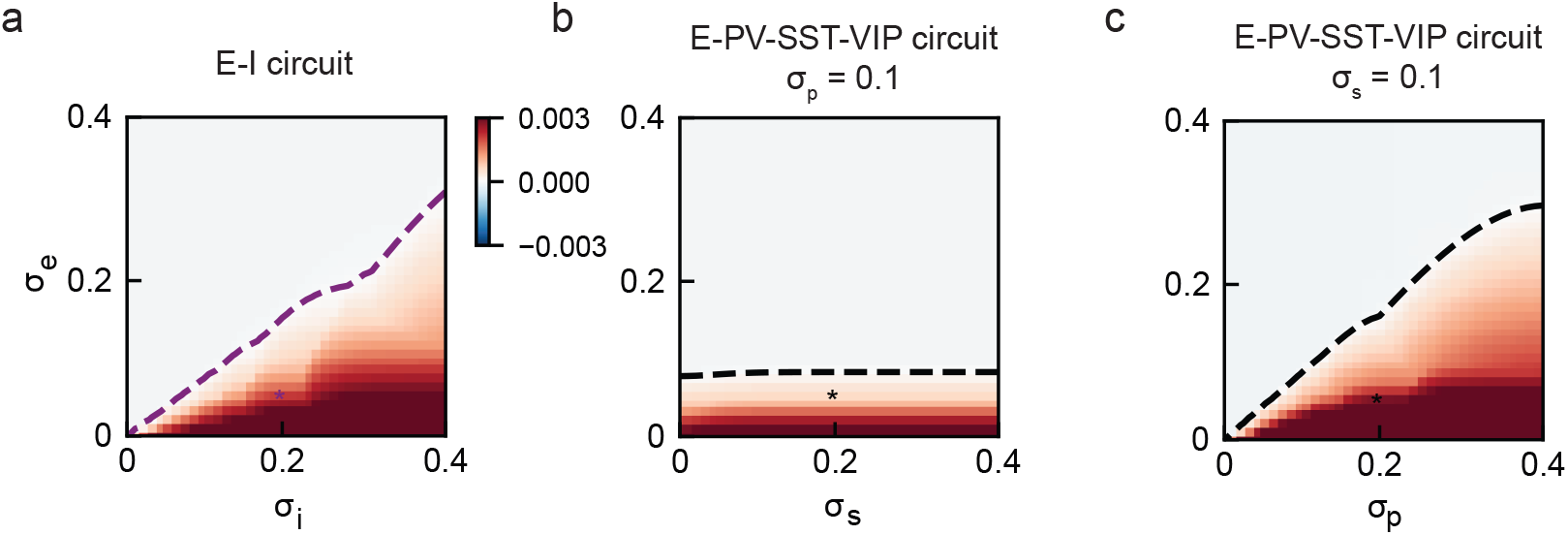
Stability diagram with *N* = 10^8^ for (a) the circuit with homogeneous inhibitory neurons, (b) the heterogeneous circuit on the *σ*_*e*_ − *σ*_*s*_ plane (*σ*_*p*_ = 0.1 fixed), and (c) the heterogeneous circuit on the *σ*_*e*_ − *σ*_*p*_ plane (*σ*_*s*_ = 0.1 fixed).

**Figure S2:**
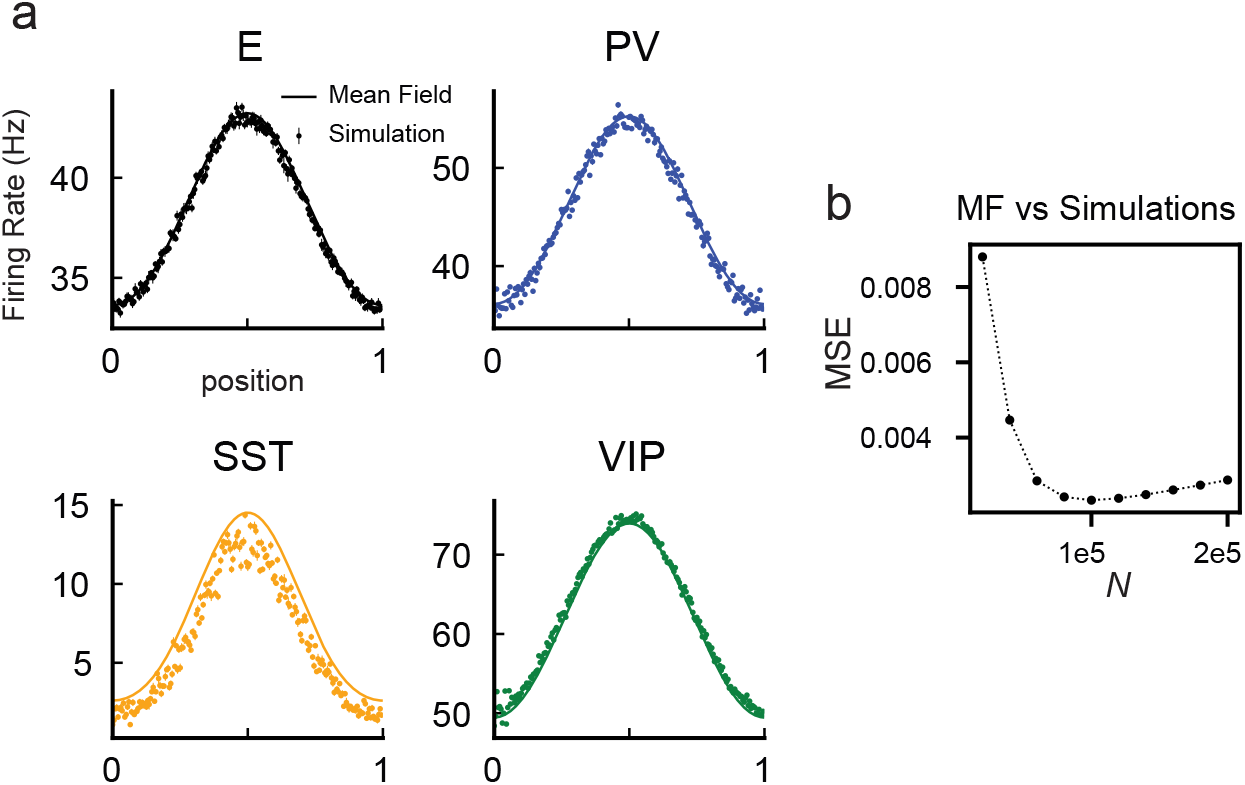
Firing rates in the four-population model with conductance-based synapses. (a) Stationary firing rate from mean field theory (solid lines) and firing rates calculated from simulations (error bars corresponding to mean ± sem; 20 independent simulations) with spiking networks of *N* = 5 × 10^5^. (b) Mean square error (MSE) between mean field theory and simulation firing rates as a function of network size *N* .

**Figure S3:**
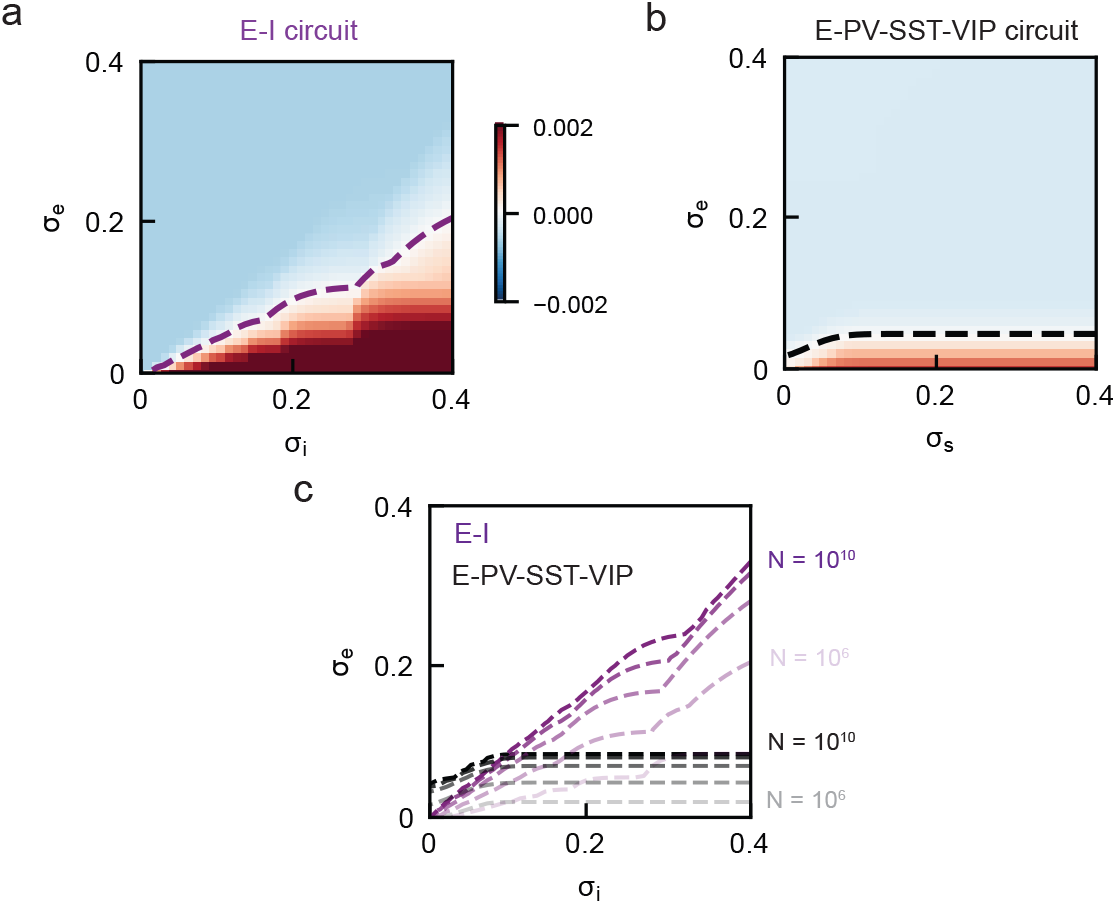
Stability with conductance-based synapses. (a) E and homogeneous I stability diagram with *N* = 10^7^ (b) Stability with the E-PV-SST-VIP circuit (c) Stability boundaries with varying network size *N* for the 2-population (black) and 4-population (purple) models.

### 1. Analysis of SST projections and stability

The stability boundary is defined by the parameters for which det *C*(*n*^*∗*^) = 0 for some critical *n*^*∗*^, where 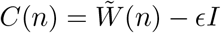 . Taking the Schur complement, we can write the determinant as

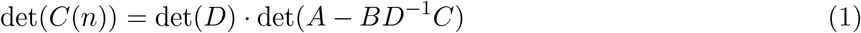

where

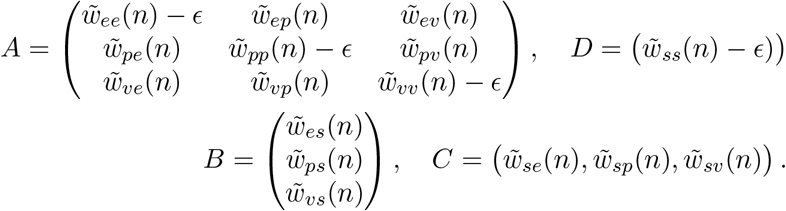

Eq. 1 indicates that the critical *n*^*∗*^ may emerge by either satisfying det(*D*) = 0 or det(*A* − *BD*^*−*1^*C*) = 0. Examining *D*, we see that 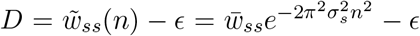. Because 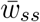 is small, the critical *n*^*∗*^ does not emerge through det(*D*) = 0: for *N* = 300000, which we use for simulations, 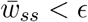 so *D* is unable to induce the bifurcation. Instead, the dependency of *σ*_*s*_ lies in the second term of *A* − *BD*^*−*1^*C*. Noting that 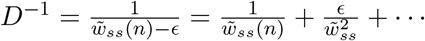, the components of *BD*^*−*1^*C* have the form

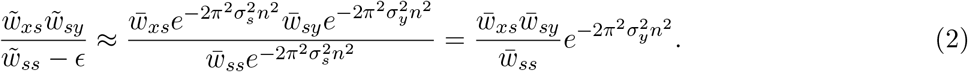

to first order in 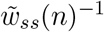. So to first order, the stability boundary is independent of *σ*_*s*_, with *σ*_*s*_-dependence emerging only when *σ*_*s*_ is very small. This is in contrast to the E-I circuit, which was already analyzed by Rosenbaum and Doiron (2014), the system becomes unstable roughly when *σ*_*e*_ ≈ *σ*_*i*_ for large *N* (assuming other conditions for stability are met).

The Schur decomposition can be interpreted as follows: there stabilization by an inhibitory population can occur through a self-dampening represented by *D* and through the rest of the network via *A* − *BD*^*−*1^*C*. However, spatial projections only have a significant effect through *D*. Therefore, if an inhibitory neuron is not stabilizing the network through self-inhibition, stability-breaking bifurcation will not occur by changing its spatial projections.

PV neurons in the E-PV-SST-VIP circuit have strong self-inhibition. To examine the effects of *σ*_*p*_ on the E-PV-SST-VIP circuit, we may repeat the analysis above but now take the Schur complement with respect to the PV component. Then we have 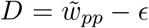. In the PV case, due to strong self-inhibition, we have 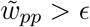 and increasing *σ*_*p*_ will result in destabilization through the factor *D*. This can be verified by repeating the analysis of Fig 3(a) but on the *σ*_*e*_ − *σ*_*p*_ plane with fixed *σ*_*s*_, in which we see a stability boundary strongly dependent on *σ*_*p*_, resembling the e-i circuit.

### 2. Parameters used in gain modulation simulations

Synaptic weights and spontaneous input current parameters used for gain modulation results in Fig. 5 are listed in Table 1. The modulatory signal 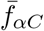 to specific inhibitory populations was given by 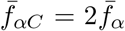 under all conditions for the current-based model; this strength was sufficient to cause partial suppression of at least one population in the network, allowing nonlinear effects. For the conductance-based model, they were set to 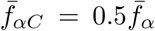 for distal stimulations. For proximal stimulations, the modulations had to be reduced further to prevent complete suppression, resulting in stimulation strenghts of 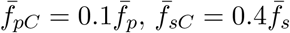, and 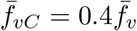.

**Table 1:**
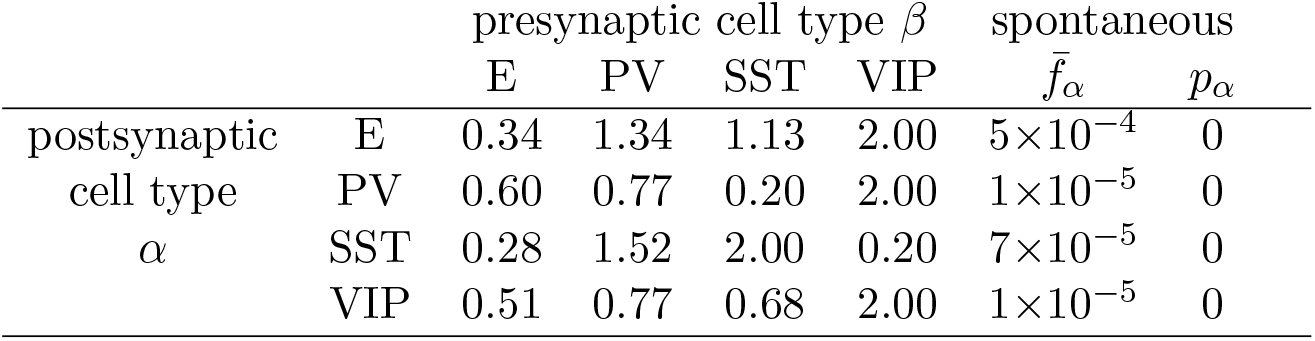
Synaptic weights *j*_*αβ*_ and spontaneous current input parameters 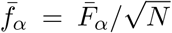 and *p*_*α*_ used in the gain modulation results in Fig. 5. Values of *j*_*αβ*_ and 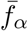 were chosen to achieve balance in the thermodynamic limit as described in the Methods. The values used for the visual stimulus parameters 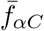 depend on the model and are listed in the main text.

| | | presynaptic cell type $\beta$ | | | | spontaneous | |
| --- | --- | --- | --- | --- | --- | --- | --- |
| | | E | PV | SST | VIP | $\bar{f}_\alpha$ | $p_\alpha$ |
| postsynaptic<br>cell type<br>$\alpha$ | E | 0.34 | 1.34 | 1.13 | 2.00 | $5 \times 10^{-4}$ | 0 |
| | PV | 0.60 | 0.77 | 0.20 | 2.00 | $1 \times 10^{-5}$ | 0 |
| | SST | 0.28 | 1.52 | 2.00 | 0.20 | $7 \times 10^{-5}$ | 0 |
| | VIP | 0.51 | 0.77 | 0.68 | 2.00 | $1 \times 10^{-5}$ | 0 |

**Figure S4:**
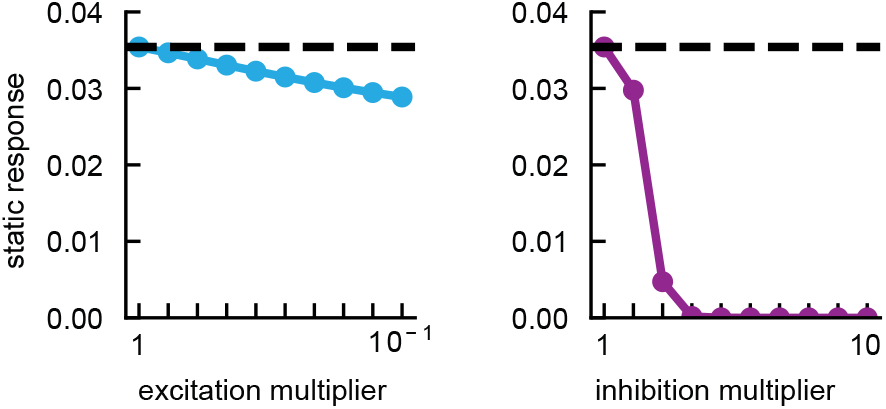
Linear response of excitatory neurons to static perturbations when excitation is decreased (left panel) and when inhibition is increased (right panel). The dashed line indicates the response when excitation and inhibition is in control network conditions.

